# Ambient Temperature Bacterial Large Ribosomal Subunit Structure Enabled by Serial Femtosecond X-ray Crystallography

**DOI:** 10.1101/2023.10.24.563633

**Authors:** Bilge Tosun, Yashas Rao, Ebru Destan, E. Han Dao, Fatma Betul Ertem, Merve Yilmaz, Ilkin Yapici, Mehmet Gul, Esra Ayan, Jerome Johnson, Alaleh Shafiei, Cahine Kulakman, Brandon Hayes, Mengning Liang, Chun Hong Yoon, Zhen Su, Mark S. Hunter, Christopher Kupitz, Frederic Poitevin, Halil Ibrahim Ciftci, Raymond G. Sierra, Steven T. Gregory, Pohl Milon, Ozge Kurkcuoglu, Hasan DeMirci

## Abstract

Ribosomes are the supramolecular complexes responsible for protein synthesis. The large 50S ribosomal subunit catalyzes the peptidyl transferase reaction and peptide bond formation between amino acids. The 50S is targeted by many known clinically effective antibiotics. Available structures, obtained at cryogenic temperatures (CT), are used for drug discovery despite that active or important target sites may display a structural configuration that is CT-induced. The introduction of ultrafast and ultrabright X-ray free electron laser (XFEL) pulses has enabled the structural observation of biological macro- and supramolecules at previously unattainable, near-physiological temperatures. In this study, we use ultrafast and ultrabright XFEL pulses to solve the apo form of 50S ribosomal subunit isolated from the extremely thermophilic bacterium *Thermus thermophilus* at ambient temperature (AT). The dimeric structure of the 50S subunit presented in this work is among the largest (∼3 megadalton) structures determined using an XFEL source to date. This study demonstrates the ability to obtain new information about ribosome structural dynamics at AT through serial femtosecond X-ray crystallography (SFX). This allowed us to capture previously unobserved dynamics of ribosomal protein uL23 and coordination by hexahydrated magnesium cations at a *hitherto* unseen resolution at near-physiological temperature. Also, residue A2602, at the core of the peptidyl transferase center (PTC), shows a rather different orientation of the sugar moiety if compared to CT structures. In addition, our structure highlights the importance of flexible residues at both the PTC and in the binding sites for antibiotics erythromycin and chloramphenicol. The method implemented here may also serve as a starting point for future structural research involving the 50S subunit complexes by employing time-resolved mix-inject and probe kineto-crystallography experiments at XFELs. Unveiling ligand-dependent 50S dynamics at physiological temperatures shall guide further development of next-generation antibiotics that target the translation machinery.

## INTRODUCTION

Ribosomes are the cellular supramolecular machineries responsible for protein synthesis (Green & Noller, 1997). The complete bacterial 70S ribosome is composed of two subunits (Steitz & Moore, 2003). The small 30S ribosomal subunit contains the decoding center which deciphers the codons on the messenger RNA (mRNA) to select the proper aminoacyl-transfer RNAs (aa-tRNAs) (Ogle et al., 2001). The large 50S ribosomal subunit contains the peptidyl transferase center (PTC) which catalyzes peptide bond formation between the amino acids attached to the A- and P-site tRNAs before translocation of the nascent peptide chain towards the peptide exit tunnel (Bashan et al., 2003). Ribosomes isolated from the extremely thermophilic bacterium *Thermus thermophilus (T. thermophilus)* have been consistently used as a model system since the earliest diffraction patterns of the 50S and 30S subunits and 70S ribosomes (Volkmann et al., 1990; Trakhanov et al., 1987). As *T. thermophilus* carries only two copies of rRNA genes, it has a significantly reduced mutational heterogeneity and also displays greater sensitivity to a broad range of antibiotics (Carr et al., 2015). These factors collectively make *T. thermophilus* an ideal target for mutational and functional analysis of ribosomes at high resolution (Gregory et al., 2005).

The *T. thermophilus* 30S small ribosomal subunit and 70S ribosomes and their complexes have been extensively studied by many groups at high-resolution at cryogenic temperatures (Yonath et al., 1988; Bulkley et al., 2012; Santos et al., 2013; Polikanov et al., 2015). Crystals of the *T. thermophilus* 50S ribosomal subunit, diffracting to 9.0 Å resolution were obtained in 1990s, leading to preliminary crystallographic studies (Volkmann et al., 1990). Furthermore, in 1991, crystals of *T thermophilus* 70S ribosome with oligo(U) and Phe-tRNA^Phe^ were obtained and diffraction data was collected up to 15 Å from a single crystal at cryo-temperature (Yusupova et al., 1991). The delay in reports of high-resolution structures of ribosomes was attributed to the intrinsic nature of traditional X-ray crystallography due to radiation damage (Volkmann et al., 1990),(Murray & Garman, 2002). Although high-resolution structures of the 50S subunit were obtained from both archeal and other bacterial sources, obtaining a crystal structure of the isolated 50S subunit of *T. thermophilus* has remained as a crystallographic challenge (Ban et al., 2000; Harms et al., 2001).

The introduction of serial femtosecond crystallography (SFX), conducted with XFELs is an alternative robust method that mitigates the radiation damage problem (Johansson et al., 2017). One significant difference between SFX and traditional synchrotron cryo X-ray crystallography is the implementation of the ‘diffraction-before-destruction’ approach, essentially collecting diffraction data before any radiation damage can occur (Neutze et al., 2000). Although the SFX method requires a large number of crystals; micro- and nano-sized crystals may be used, allowing data collection from complex biological macromolecules like ribosomes without growing large crystals (DeMirci et al., 2013). Additionally, SFX does not require the cryo-cooling process employed in synchrotron cryo-crystallography which, despite reducing crystal damage from radiation, may induce crystal damage by mechanical shearing <u>(Sierra et al., 2016)</u>. These differences have enabled structural data collection at ambient temperature with minimal structural/lattice damage (Boutet et al., 2012).

Here we report the ambient-temperature dimeric apo-form of large 50S ribosomal subunit structure determined at the Coherent X-ray Imaging (CXI) instrument of the Linac Coherent Light Source (LCLS) (Boutet et al 2012), displaying 1712 associated well-defined hexahydrated magnesium cations dispersed throughout the unit cell beyond 4 Å resolution. To emphasize the effect of temperature on ribosomal conformation, we compared our structure with previous cryogenic-temperature structures through superposition and Gaussian Network Modelling (GNM) analysis. Our structure and GNM analysis reveals several significant conformational changes in functionally-important 50S ribosomal protein and rRNA residues. Collectively these proof-of-the-concept data demonstrate the feasibility of obtaining near-physiologic temperature SFX structures which will enable further future structural studies of antibiotics targeting the large 50S ribosomal subunit and also 70S ribosome functional complexes.

## MATERIAL AND METHODS

### Purification and crystallization of 50S ribosomal subunits

Growth of *T. thermophilus* HB8 strain was performed as previously described (DeMirci, Murphy, et al., 2013; DeMirci et al., 2010) using ATCC:697 *Thermus* medium enriched with Castenholz salts at 72 ℃ under intense aeration. The cells were grown in 4 L baffled glass flasks by using New Brunswick Innova 4430 incubator shaker and harvested at OD_600_=0.8 by centrifugation at 4000 rpm at 4℃. Cell pellets were cooled by flash freezing in liquid nitrogen and stored at −80 ℃. To resuspend the cells, Buffer A containing 100 mM ammonium chloride (NH_4_Cl), 10.5 mM magnesium acetate, 0.5 mM Ethylenediaminetetraacetic acid (EDTA), 20 mM N-2-hydroxyethylpiperazine-N-2-ethane sulfonic acid (HEPES), adjusted with potassium hydroxide (KOH) to pH 7.5 was used. Resuspended cells were disrupted by two passes through an emulsiflex C5 (Avestin, ON, Canada) initially at 12,000 psi and second pass at 20000 psi. As the next step, the samples were centrifuged at 20000 rpm for 20 minutes with a Ti-45 rotor (Beckman Coulter, USA) to remove the cell debris. Supernatants of the sample applied to sucrose cushion Buffer B containing 1.0 M NH_4_Cl, 1.1 M sucrose, 10.5 mM magnesium acetate, 0.5 mM EDTA, 20 mM HEPES-KOH (pH 7.5) were centrifuged at 43000 rpm for 15 hours with a Ti-45 rotor. After rewashing the samples with Buffer A, the pellets were resuspended with Buffer C containing 1.5 M ammonium sulfate (NH_4_)_2_SO_4_, 10 mM magnesium acetate (MgOAc), 20 mM HEPES-KOH (pH 7.5). After filtration of the resuspended ribosomes, the sample was loaded onto the POROS-ET hydrophobic interaction column equilibrated with Buffer C (TOSOH Japan). Afterwards, reverse ammonium sulfate gradient containing Buffer C and Buffer D containing 20 mM HEPES-KOH (pH 7.5) and 10 mM MgOAc was used for elution of the samples and 70S peak fractions were collected. The collected fractions pooled and were dialyzed into Buffer E containing 50 mM KCl, 10 mM NH_4_Cl, 10 mM HEPES-KOH (pH 7.5), 10.25 mM MgOAc, 0.25 mM EDTA. To concentrate the fractions, Amicon 30 KDa PM30 an ultrafiltration membrane was employed. Separation of the 30S and 50S subunit fractions containing a mixture of both subunits was achieved by applying 10%-40% linear sucrose gradient in Buffer F containing (50 mM KCl, 10 mM NH_4_Cl, 10 mM HEPES-KOH (pH 7.5), 2.25 mM MgOAc, 0.25 mM EDTA. Then the zonal ultracentrifugation was performed for 16 hours at 27000 rpm in a Beckman Ti-15 rotor. Fractions were recovered from the top of the gradient by pumping a 55% sucrose solution to the bottom of the Ti-15 zonal rotor. Then 50S peak was dialyzed into Buffer C for further separation by the POROS-ET column with reverse ammonium sulfate gradient. RNAse containing fractions were discarded to extend the stability of the large 50S ribosomal subunits during crystallization and the remaining RNase-free fractions were combined and dialyzed into Buffer G containing 50 mM KCl, 10 mM NH_4_Cl, 5 mM HEPES-KOH (pH 7.5), 10 mM MgOAc and to be concentrated by Amicon 30 KDa PM30 an ultrafiltration membrane to final OD_260_=400.

The apo 50S subunits were screened for crystallization by using 3500 commercially available grid and sparse matrix crystallization cocktails performed under sitting-drop microbatch screening technique under oil (DeMirci, Murphy, et al., 2013; Dao et al., 2018; O’Sullivan et al., 2018). We obtained more than 20 conditions yielding SFX quality crystals (Supplementary Figure 1). The best selected crystallization condition contained 200 mM ammonium citrate tribasic pH 7.0, 20% w/v PEG 3350 solution. Microseeding and batching the 1 ml ribosome samples 1:1 v/v ratio yielded *T. thermophilus* 50S crystals of approximate dimensions 3-5 μm × 3-5 μm × 5-10 μm (Supplementary Figure 1). Crystal concentration was approximated to be 10^9^–10^10^ particles per ml based on light microscopy and nanoparticle tracking analysis NanoSight LM10-HS with corresponding Nanoparticle Tracking Analysis (NTA) software suite (Malvern Instruments, Malvern, UK).

### Sample Delivery with coMESH

Data collection of microcrystals at XFEL requires gentle and robust sample-delivery methods specifically accommodating the ‘diffract-before-destroy’ approach and the femtosecond pulse-length of XFEL sources (Martiel et al., 2019). For ultra-delicate 50S ribosome crystalline sample delivery the selected approach was the concentric Microfluidic Electrokinetic Sample Holder (coMESH) due to its elimination of inline filters and low sample consumption <u>(Sierra et al., 2016)</u>. The coMESH injector has been used successfully for many ribosome XFEL data collection experiments and it uses coaxial internal capillaries to deliver two solutions via an ‘inner’ sample capillary and an ‘outer’ sheet capillary <u>(Sierra et al., 2016</u>; Dao et al., 2018; O’Sullivan et al., 2018). This approach was selected due to favorable hit-rates and resultant data quality observed with prior macromolecules probed at XFELs <u>(Sierra et al., 2016</u>; Dao et al., 2018; O’Sullivan et al., 2018). The crystalline samples suspended in unaltered mother liquor were delivered to the interaction region of the XFEL beam via the inner concentric sample capillary. A ‘sister liquor’, delivered co-axially and simultaneously via a larger-diameter outer capillary, prevents sample freezing and dehydration in the vacuum environment of the CXI instrument of the LCLS.

Initially the sister liquor was loaded, flowed and electrically focused. After obtaining a slightly stable jet, the central sample line was connected. The air coming from the previously disconnected sample line introduces bubbles, so the sister liquor cannot be stabilized completely <u>(Sierra et al., 2016)</u>. The outer line must be operating in vacuum to prevent jet blockages and cloggings, since it has noticeably less fluidic resistance compared to the central sample line. The sister liquor’s flow rate and the mother liquor’s flow rate are equal to each other to be 1 uL/min, to ensure a stable injector and to expose maximum number of crystals to the X-ray interacting region (O’Sullivan et al., 2018).

The sample reservoir was loaded with the 50S ribosome crystal slurry in the unaltered mother liquor containing 200 mM ammonium citrate tribasic (pH 7.0) and 20% w/v PEG 3,350. The sister liquor was similar to the mother liquor with the exception of an additional 20% MPD (v/v) concentration. The sample capillary was a 100 μm ID × 160 μm OD × 1.5 m long fused silica capillary. The sheath flow capillary was a 75 μm ID × 360 μm OD × 1 m long fused silica capillary connected to the side port of the T-junction to flow the cryoprotectant sheath liquid line. The continuous outer concentric capillary was a 200 μm ID × 360 μm OD × 5 cm long tapered silica capillary. The tips of the inner and outer capillaries were located co-terminally at the same position <u>(Sierra et al., 2016</u>). The applied voltage on the sheath liquid was typically 3000 V, and the counter electrode was grounded. The sample ran typically between 0.75 and 1 μl/min, and the sheath flow rate typically matched the sample flow or was slightly faster.

### Data collection and analysis for SFX studies at LCLS

SFX data acquisition was carried out at ambient temperature at the Coherent X-ray Imaging (CXI) instrument of the Linac Coherent Light Source (beamtime ID: cxim7916) at the SLAC National Accelerator Laboratory (Menlo Park, CA). An X-ray beam with a vertically polarized pulse of 30–40 fs duration was focused with refractive Beryllium compound lenses with a ∼6 × 6 µm^2^ full width at half maximum beam size. The experiments were performed with 9.5 keV photon energy and 120 Hz repetition rate. Real-time data analysis was performed to determine the initial diffraction geometry, monitor the crystal hit rates, and analyze the gain switching modes of the CSPAD detector (Kameshima et al., 2014) using OM monitor version 1.0 (Mariani et al., 2016) and PSOCAKE version 1.0.8 (Damiani et al., 2016; Thayer et al., 2017). A total of 337.560 detector frames were collected in 47 min continuously with the apo *T. thermophilus* 50S microcrystals. 50S microcrystals were delivered to the X-ray interacting region using coMESH (Sierra et al., 2015). Individual diffraction pattern hits were defined as frames containing more than 30 Bragg peaks with a minimum signal-to-noise ratio larger than 4.5, for a total of 38400. The detector distance was set to 210 mm, with an achievable resolution of 3.08 Å at the edge of the detector (2.6 Å in the corner). The structure of the 50S ribosomal subunit was determined at 3.99 Å resolution.

### Ambient temperature structure determination and refinement

The apo form of 50S ribosomal subunit dimer structure was determined from microcrystals in space group P4_1_. Molecular replacement was performed with the molecular replacement program *PHASER 2.8.3* (McCoy et al., 2007) within the *PHENIX version 1.19.2* software suite (Adams et al., 2010), using only the 50S large subunit part of the previously published 70S ribosome structure as an initial search model (PDB ID: 4V8I) (Polikanov et al., 2012). Using the *PHENIX* software suite, the initial rigid body refinement was performed with the 4V8I 50S large subunit coordinates. Individual coordinates and TLS parameters were refined following the simulated-annealing refinement. Potential positions of altered side chains and water molecules were identified by performing composite omit map refinement within *PHENIX*. The density map was analyzed using *COOT version 0.8.9.2* (Emsley & Cowtan, 2004). Positions with strong difference density were retained while the hexahydrated magnesium molecules outside the electron density were manually removed. The Ramachandran statistics for the 50S dimer (favored / allowed / outliers) were 96.88 / 3.07 / 0.05 %, respectively. The structure refinement statistics are shown in the crystallography table. Structural alignments were performed using *PyMOL version 2.3* (<u>DeLano, W.L. (2002) The PyMOL Molecular Graphics System. Delano Scientific, San Carlos.</u>) with the default 2σ rejection criterion and five iterative alignment cycles. All figures were generated using *PyMOL*.

### Gaussian Network Model (GNM)

The low-frequency motions of the ribosome large subunit 50S and the ribosomal protein uL23 alone were investigated using GNM. The 3-dimensional structure was described as an elastic network formed of *N* nodes (nucleotides and/or amino acids) that are connected by uniform springs if they are within a cutoff distance. Nodes were placed to P (phosphate) atoms (nucleotides) and Cα atoms (amino acids), i.e. one interaction site per residue. Three different cutoff distances were used to construct the elastic network: 10 Å to connect Cα – Cα pairs, 15 Å for Cα – P pairs, and 20 Å for P-P atom pairs (Kurkcuoglu et al., 2020). The potential energy in GNM is given based on pairwise connectivity as

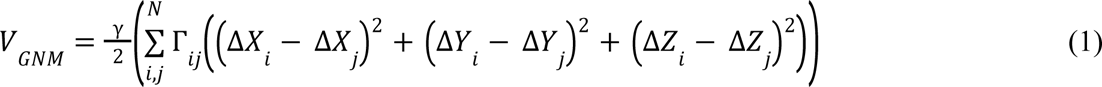

where, γ is the uniform spring constant. Δ shows the fluctuation of nodes *i* and *j*, which are assumed to be isotropic and obey Gaussian distribution (Bahar et al., 1997; Haliloglu et al., 1997). *Г_ij_* is the *ij^th^* element of the Kirchhoff matrix (*N* × *N*) showing the connectivity of the nodes. After the singular value decomposition of the Kirchhoff matrix, *N*-1 non-zero eigenvalues **λ_k_** and corresponding eigenvectors **u_k_** were obtained to describe the dynamic behavior of the nodes in *k^th^* mode of motion. Cross-correlation between the fluctuations of *i^th^* and *j^th^* nodes averaged over *k* modes was calculated as,

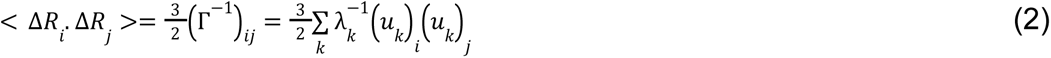

In this study, we are interested in the fluctuations at the lowest frequency of the dynamic spectrum. < Δ*R_j_*. Δ*R_j_*> Values calculated for the modes of motion of the structure reveal the distinct dynamic domains separated by flexible hinges, essential for its functional dynamics. The first three low-frequency modes (slow modes) of the bacterial ribosome structure correspond to its functional collective motions, such as ratchet-like rotation, anti-correlated motions of the L1 and bL12 stalks, and rotation of the small subunit head (Kurkcuoglu et al., 2008).

### Mg^2+^/Mg(H_2_O)_6_^2+^ and 50S ribosomal subunit interactions

Electrostatic and hydrogen bonding interactions between Mg(H_2_O)_6_^2+^, Mg^2+^ ions and the 50S ribosomal subunit were calculated with Discovery Studio 4.1 Free Visualizer using default settings to determine non-bonded interactions. Histograms of these interactions were generated with MATLAB R2017a (Istanbul Technical University (ITU) academic and research license).

## RESULTS

### Ambient temperature structure of 50S ribosomal subunit enabled by serial femtosecond X-ray crystallography

The ambient-temperature structure of the ligand-free *T. thermophilus* 50S ribosomal subunit (from here on, 50S^AT^) was determined to a resolution of 3.99 Å at the Linac Coherent Light Source (LCLS) using a low-flow concentric electrokinetic liquid injector setup (Figure 1A-B & Supplementary Table 1). The 1.5 MDa macromolecule is found as a dimeric form in the crystal lattice, resulting in a large ∼3 MDa complex packed in the asymmetric unit cell with dimensions of a=303.8, b=303.8 and c=434.8 Å in P4_1_ space group. The 47 minutes of data collection consumed approximately 50 µL of *T. thermophilus* 50S crystal sample slurry (Supplementary Figure 1). A clear electron density map was observed for the 23S and 5S rRNAs, as well as for 35 ribosomal proteins. The primary area of interest is the peptidyl transferase center (PTC) located within the core of the 23S rRNA appears well-defined in the experimental electron density map (Supplementary Figure 2). Additionally, the 50S^AT^ structure shows a dynamically stable core and flexible regions around the 5S and 23S rRNAs (Figure 1C-D). As observed for 50S structures from *Deinococcus radiodurans* and *Haloarcula marismortui*, proteins interacting with GTP-bound translation factors, including uL1, uL10, uL11, bL12 are found as disordered components of the electron density map (Ban et al., 2014); (Ban et al., 2000).

**Figure 1.**
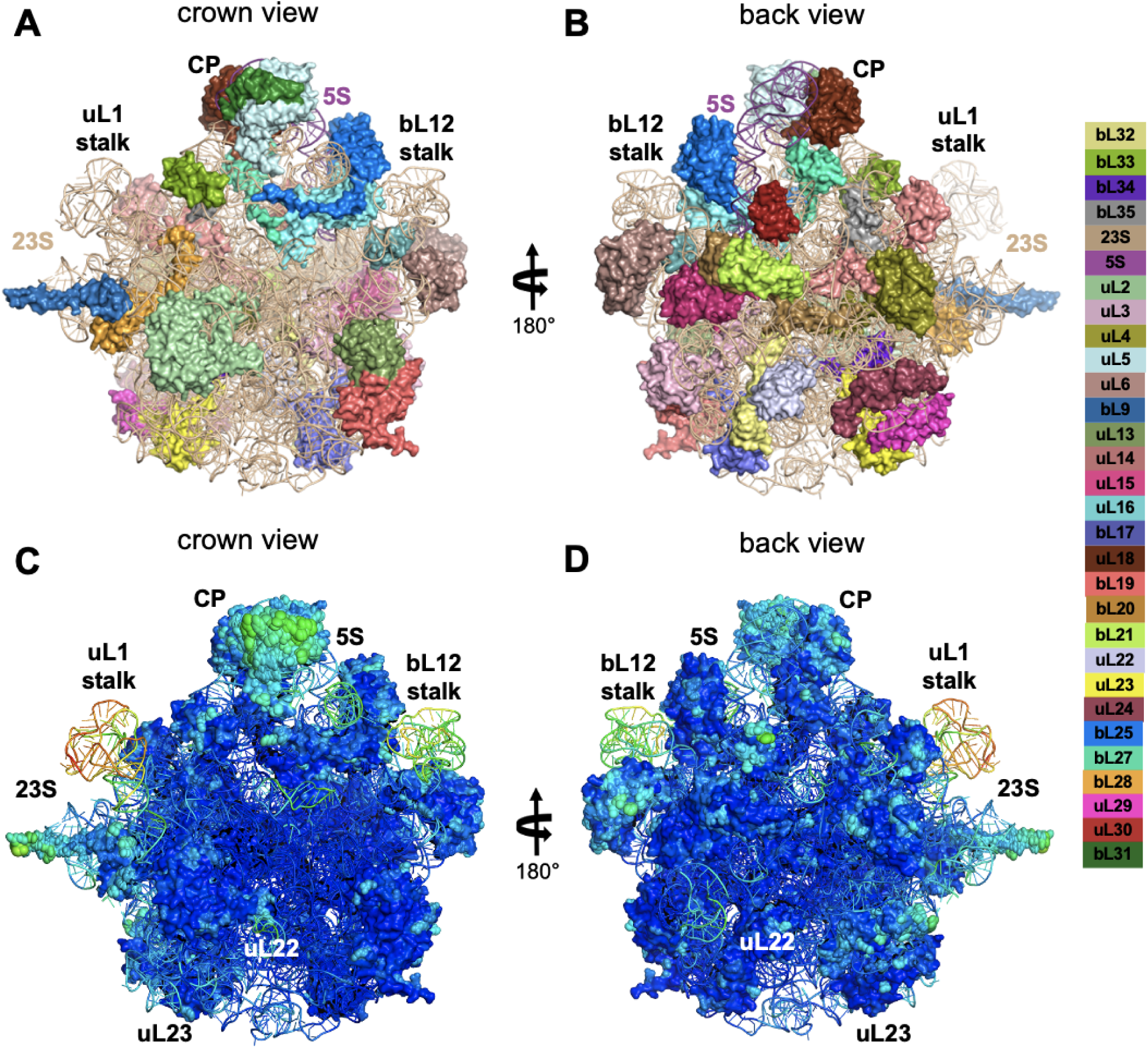
Representation of 50S ribosomal subunit from *T. Thermophilus* at ambient temperature. **(A-B)** Proteins around the 23S and 5S RNA are indicated with different colors (Ban et al., 2014). **(C-D)** Overall structure is shown with ellipsoid representation to reveal stable/flexible regions for 50S ribosomal subunit from crown view (panel A and C). The bL12 stalk is on the right, the uL1 stalk is on the left, and the central protuberance (CP) including the 5S rRNA (purple) is in the middle of the upper part of the particle.

The 50S^AT^ dimeric crystal structure reveals the interactions between the individual large ribosomal subunits for the first time (Figure 2A). Dimerization in our structure is mediated by the interaction between the 23S RNA and the L1 stalk (Figure 2B-C). In the 50S^AT^ structure, the stalk is stabilized in the crystal lattice by being involved in dimeric packing (Figure 2B-C). Residues A2180, G2181, C2183, G2184, C2188, G2190, A2191, and A2192 are part of the linker site that connect both 50S subunits, and the residues interact with the surrounding Mg^2+^ cations. Additionally, bL33 residues Pro31, Asn32, and Lys33 form hydrogen bonds with A2186, G2187, and G2188, from the opposite 50S subunit, likely stabilizing the interaction (Figure 2B). These interactions are anticipated to play a crucial role for the dimeric packing process, particularly for the residues that are observed to be flexible in the linker site (Figure 2C). Altogether, the dimeric structure of the 50S^AT^ is among the largest (∼3 megadalton) structures determined using an XFEL source to date in addition to be the first crystals of the *T. thermophilus* 50S ribosomal subunit solved in the absence of other ligands.

**Figure 2.**
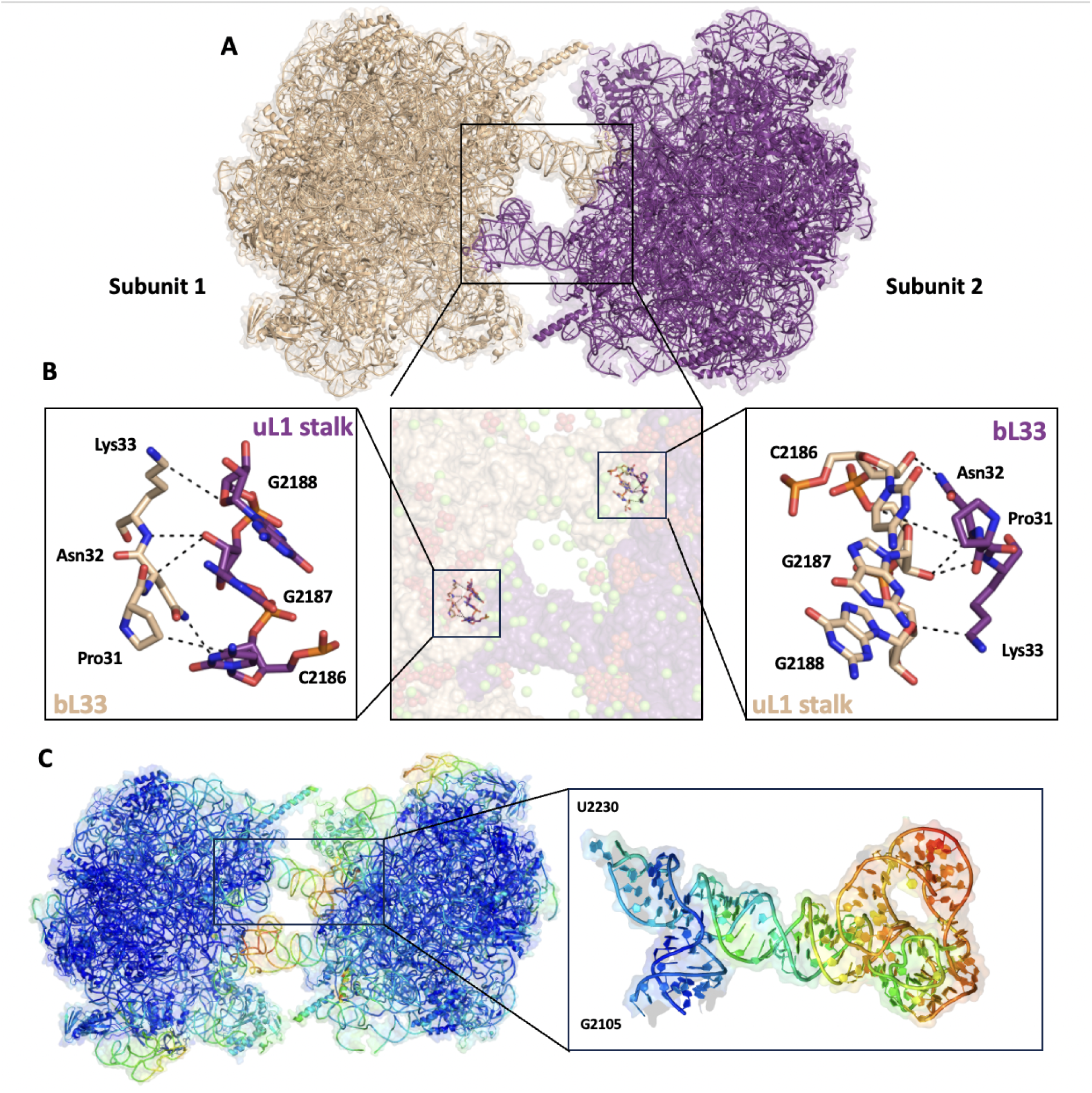
Dimeric 50S ribosomal subunit structure in asymmetric unit cell. **(A)** Overall structure of dimeric 50S ribosomal subunit (subunit 1 colored as wheat and subunit 2 colored as purple) in cartoon representation. (**B)** Close up view of dimer surfaces; the mid panel shows the subunits as surfaces, Mg^2+^ in green spheres and hexahydrated Mg^2+^ cations in green and red spheres. Left side panel shows the interactions between bL33 of subunit 1 and uL1 stalk of subunit 2; the right side panel shows the interactions between uL1 stalk of subunit 1 and bL33 of subunit 2. The interactions on both sides occur through the hydrogen bonds between the residues C2186, G2187, and C2188 in uL1 stalk and Asn32, Pro31, and Lys33 of bL33 protein. **(C)** B-factor illustration of dimerization. The regions colored red are more flexible, while the regions colored blue are more rigid. 23S rRNA has flexible regions where it interacts with the bL33 protein of the other monomer.

### Increased solvent presence

Another advantage of XFELs compared to synchrotron-based light sources is that the former collects additional features of the solvent content since the solvent is highly susceptible to photoelectric absorption (Weik et al., 2000). The negatively-charged phosphate backbone of RNA requires cationic stabilization, especially within the ribosome due to the extensive RNA folding to form the compact rRNA-dense structure (Klein et al., 2004). The phosphate backbone stabilization is provided by metal cations, typically Mg^2+^, Na^+^, and K^+^, depending on the buffer used. In a magnesium-depleted medium, complete ribosomal degradation occurs (McCarthy, 1962). A comparison of the 50S^AT^ with cryogenic temperature crystal structures of (50S^CT^) highlights the broad dispersion of solvent within the crystals. The cryogenic structure of the *H. marismortui* 50S^CT^ (PDB ID: 1FFK) captures ∼120 Mg^2+^ cations stabilizing the 23S rRNA. In contrast our *T. thermophilus* 50S^AT^ structure shows experimental electron densities for 1712 hexahydrated and hundreds of stand-alone magnesium cations within and surrounding the 23S rRNA due to low photoelectric absorption and mitigated radiation damage (Figure 3).

**Figure 3.**
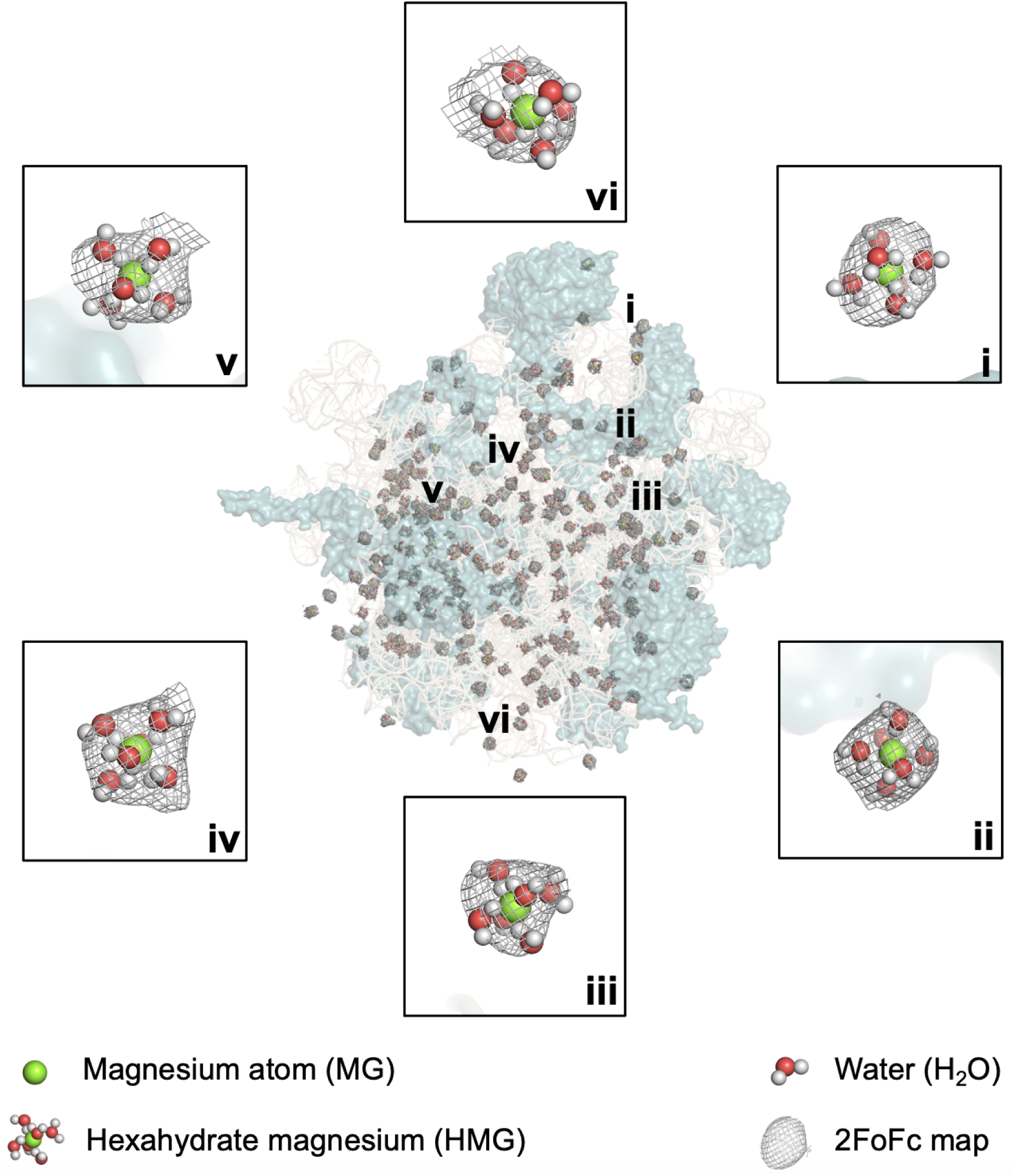
Representation of hexahydrated magnesium cations around 50S ribosomal subunit. **(A-B)** 50S ribosomal subunit is shown with cartoon representation and hexahydrate magnesium cations are shown in lightteal around the 50S ribosomal subunit. 2*Fo-Fc* simulated annealing-omit map for HMG is shown in gray and contoured at 1.0 σ level.

The hydrated Mg^2+^ cations were determined iteratively by scanning the weighted *Fo-Fc* electron-density difference map at 3σ. The resulting electron densities were visually determined to be hexahydrated magnesium ions, which have six water molecules arranged in an octahedral arrangement, whereas the smaller densities were determined as dehydrated Mg^2+^ cations. A total of 1712 hexahydrated magnesium ions were found in and around 5S and 23S rRNA and associated proteins within the 50S^AT^ dimer. For the first monomer, a total of 860 hexahydrated magnesium ions were determined. Within them, 113 form hydrogen bonds with polar surface residues of the proteins or the carbonyl group of their backbone. However, 536 hexahydrated magnesium ions interact with the 23S rRNA while 20 bind to the 5S rRNA at the phosphate backbone or the base itself. A total of 852 hexahydrate magnesium ions were found interacting with the second monomer. Within them, 192 form hydrogen bonds with polar surface residues of the proteins or the carbonyl group of their backbone. 720 hexahydrated magnesium ions bind to the 23S rRNA while 13 bind to the 5S rRNA at the phosphate backbone or the base itself.

Additionally, we performed an extensive analysis of the non-bonded interactions of hexahydrated magnesium and stand alone magnesium cations in the 50S^AT^ ribosomal subunit. Attractive charges and pi-cation interactions involving magnesium ions, as well as hydrogen bond and pi-hydrogen bond interactions of oxygen atoms are presented for the hexahydrated magnesium ions on one 50S^AT^ subunit by histograms (Supplementary Figure 3A). These groups have the capacity to establish up to 19 non-bonded interactions at the same time with different nucleotides and/or amino acids within a distance of ∼5.5 Å. The majority of these interactions are formed by hydrogen bonds, followed by electrostatic interactions. Few pi-cation and pi donor-hydrogen bond interactions were also noted. The interactions of hexahydrated magnesium ions were found to be mostly with G nucleotides of 23S rRNA and least with U nucleotides (Supplementary Figure 3B). Attractive charge interactions involve Asp, Glu and Val residues on numerous ribosomal proteins. Hydrogen bond interactions are noted to occur especially from the backbone oxygen, side chain oxygen or nitrogen atoms of the amino acids, reflected in the contribution of a high number of ribosomal proteins. A small number of pi-cation or pi donor-hydrogen bond interactions were found for aromatic amino acids His, Trp and Tyr, as expected. The binding architecture of hexahydrated magnesium ions is illustrated by 23S rRNA and ribosomal protein uL23 interface (Supplementary Figure 3C). The magnesium ion makes electrostatic interactions with three residues, namely A1388, U1648 and Thr56. Numerous hydrogen bond interactions were noted to stabilize the rRNA-protein interface, while a stable pi donor-hydrogen bond interaction also occurs with the base of 23S rRNA C1646. On the other hand, magnesium ions on the 50S subunit can establish up to 6 non-bonded interactions with rRNAs and/or ribosomal proteins (Supplementary Figure 4). Attractive charge interactions mostly involve G nucleotides of 23S and 5S rRNAs, as well as Glu and Asp amino acids of ribosomal proteins. Pi-cation interactions are also populated near G nucleotides of 23S rRNA.

The 50S^AT^ crystal structure and architecture is first described here for apo *T. thermophilus* 50S subunits, albeit a considerable number of cryogenic 70S structures are available. In order to establish a comparable framework, we first aligned our 50S^AT^ to several 50S^CT^ structures extracted from their respective 70S^CT^. Specifically, the surface and intersubunit bridging residues were compared to minimize a possible structural influence from the bound 30S^CT^. Our analysis shows a large degree of stability with few examples of Mg^2+^ ions and ribosomal residues showing different orientations. This, in turn allows us to compare the 50S^AT^ structure with, in the absence of a solved 50S^CT^ crystal, *in bona fede* the most similar 50S structures obtained at cryogenic temperatures. From now on, the comparisons will be denoted by 50S^CT^ followed by the PDB codes.

### uL23 is observed in a novel conformation

Proteins around the 50S ribosomal subunit were analyzed to detect conformational changes due to temperature. Interestingly, uL23 in the 50S^AT^ structure has an extended loop conformation (Val59 to Arg76 in *T. thermophilus*) if compared to the 50S^CT_Holo^ from *T. thermophilus* and bound to erythromycin-(PDB: 4V7X) (Figure 4A). This may result from differences on the bound drug or influenced by temperature (Figure 4A). To further inquire, we compared 50S^AT^ to the 50S^CT_Apo^ obtained in the absence of any drug (PDB ID: 4V8I) and observed uL23 displaying similar configuration at the extended loop, near the exit tunnel. However, Arg68 is positioned differently in the structures, suggesting a certain degree of flexibility (Figure 4B). To further study these conformational changes, the flexibility and functional dynamics of uL23 we used Gaussian Network Models (GNM). We first calculated the theoretical B-factors of uL23 in the 50S subunit and uncomplexed uL23 using all normal modes. Then, we compared the normalized values with the experimental B-factors at ambient temperature (Figure 5A). GNM results suggested that the mobility of Arg68 at the tip of the long loop of uL23 bound to 50S is lower than Arg68 of uncomplexed uL23. Structural data at ambient temperature also supported this finding, as Arg68 makes a salt bridge with C482 of 23S rRNA that would result in an extended conformation but relatively limited mobility. On the other hand, the stem of the extended loop is expected to be immobile based on GNM calculations for uL23 in 50S^CT^ the 50S subunit. However, the stem is rather mobile at ambient temperature similar to its uncomplexed dynamics calculated with GNM. In addition, we analyzed the functional motions of the extended loop using the low-frequency modes of GNM (first 10 slow modes) (Figure 5B-C). It was noted that the extended loop of uL23 is a distinct dynamic domain separated from the globular core of the protein at lowest-frequency motions, with or without incorporation into the 50S subunit. This would facilitate its interaction with emerging peptides through the exit tunnel. The dynamic domains shaped by the cumulative 10 slowest modes indicated different hinge regions on the loop, which are expected to appear during its functional motions (Figure 5C). Thus, the extended loop has a high capacity to adopt different conformations due to this dynamic feature.

**Figure 4.**
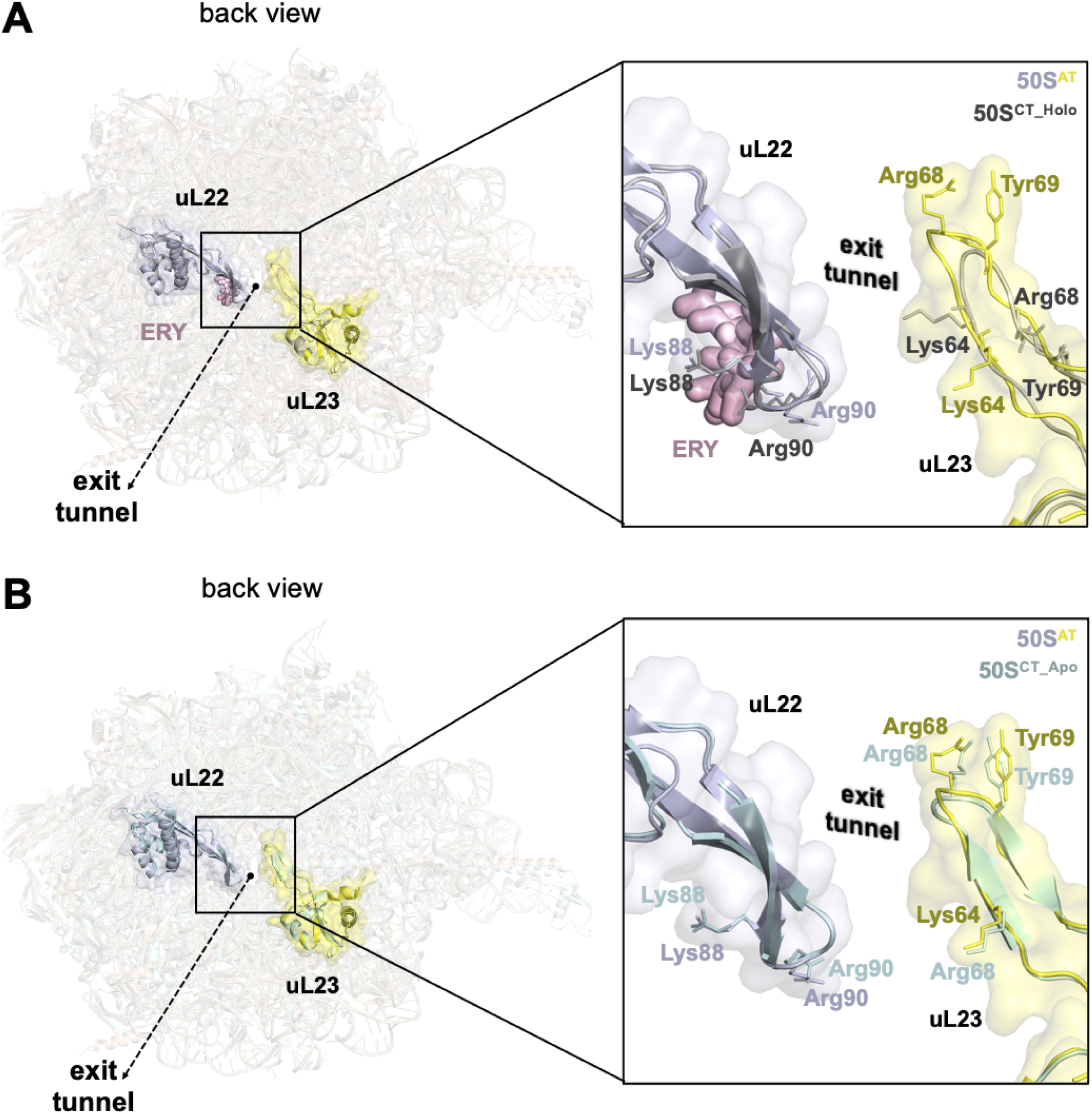
Ambient temperature dynamics of ribosomal protein uL23. **(A)** Erythromycin bound 50S structure (PDB ID: 4V7X) is superposed with the apo structure at ambient temperature to highlight the binding of drug near to exit tunnel. **(B)** Apo structure of 50S ribosomal subunit at cryogenic temperature (PDB ID: 4V8I) is superposed with the apo structure at ambient temperature to illustrate the dynamics of ribosomal protein uL23.

**Figure 5.**
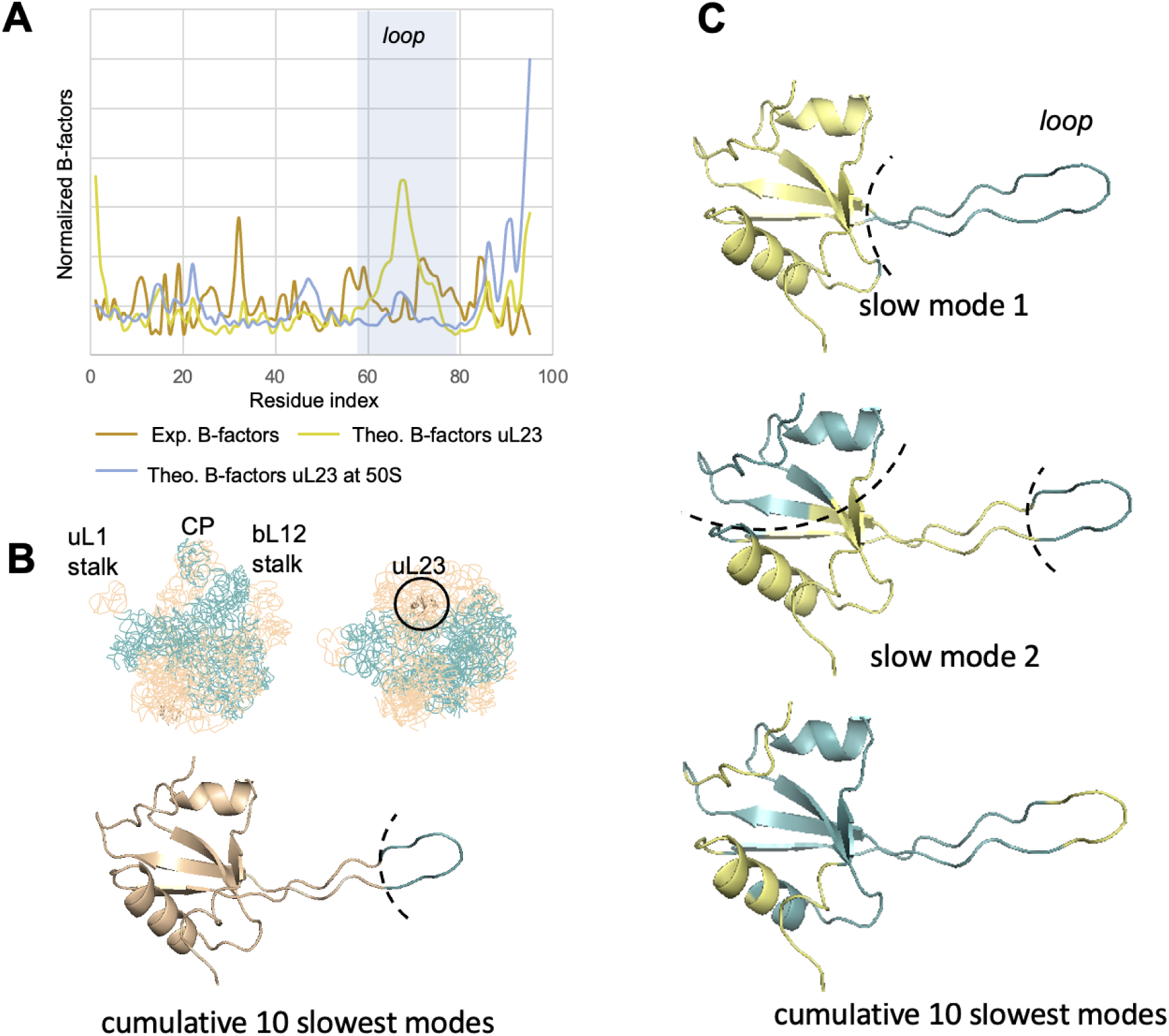
GNM analysis and functional motions of ambient temperature uL23 in the 50S ribosomal subunit. **(A)** Theoretical B-factors of uncomplexed uL23 shown in orange, uL23 in 50S shown in blue, and experimental B-factors of uL23 in 50S shown in gray. **(B)** Dynamic domains of the 50S defined by the 10 slowest modes shown from different perspectives. Dynamic domains of uL23 coordinated by the hinge region (in dashed line) are explicitly shown. (**C)** Dynamic domains shaped by slow mode 1, slow mode 2 and first 10 slow modes for the uncomplexed uL23. Dynamically correlated domains are shown with gray and blue, hinges are indicated by dashed lines in **(B)** and **(C)**.

### Peptidyl Transferase Center Residues

The peptidyl-transferase center is situated within the core of the large ribosomal subunit and catalyzes peptide bond formation. This center contains five pivotal residues, namely A2602, U2585, U2506, A2451, and C2063, positioned to facilitate the creation of peptide bonds between two amino acids (Amort et al., 2007). These residues exhibit varying degrees of flexibility, depending on their respective roles in catalytic activities. The differences in conformational states become evident when comparing the 50S^AT^ structure with superimposed 70S ribosomes and 50S ribosomal subunits from cryogenic temperature sources. Notably, our findings reveal a significant conformational alteration of the ribose sugar at the 23S rRNA residue A2602, which is crucial for efficient peptidyl-tRNA hydrolysis (Figure 6) (Polacek et al., 2003). Residue A2602, positioned between the A and P sites, can adopt various orientations and often exhibits weak electron density in crystal structures (Beringer & Rodnina, 2007).

**Figure 6.**
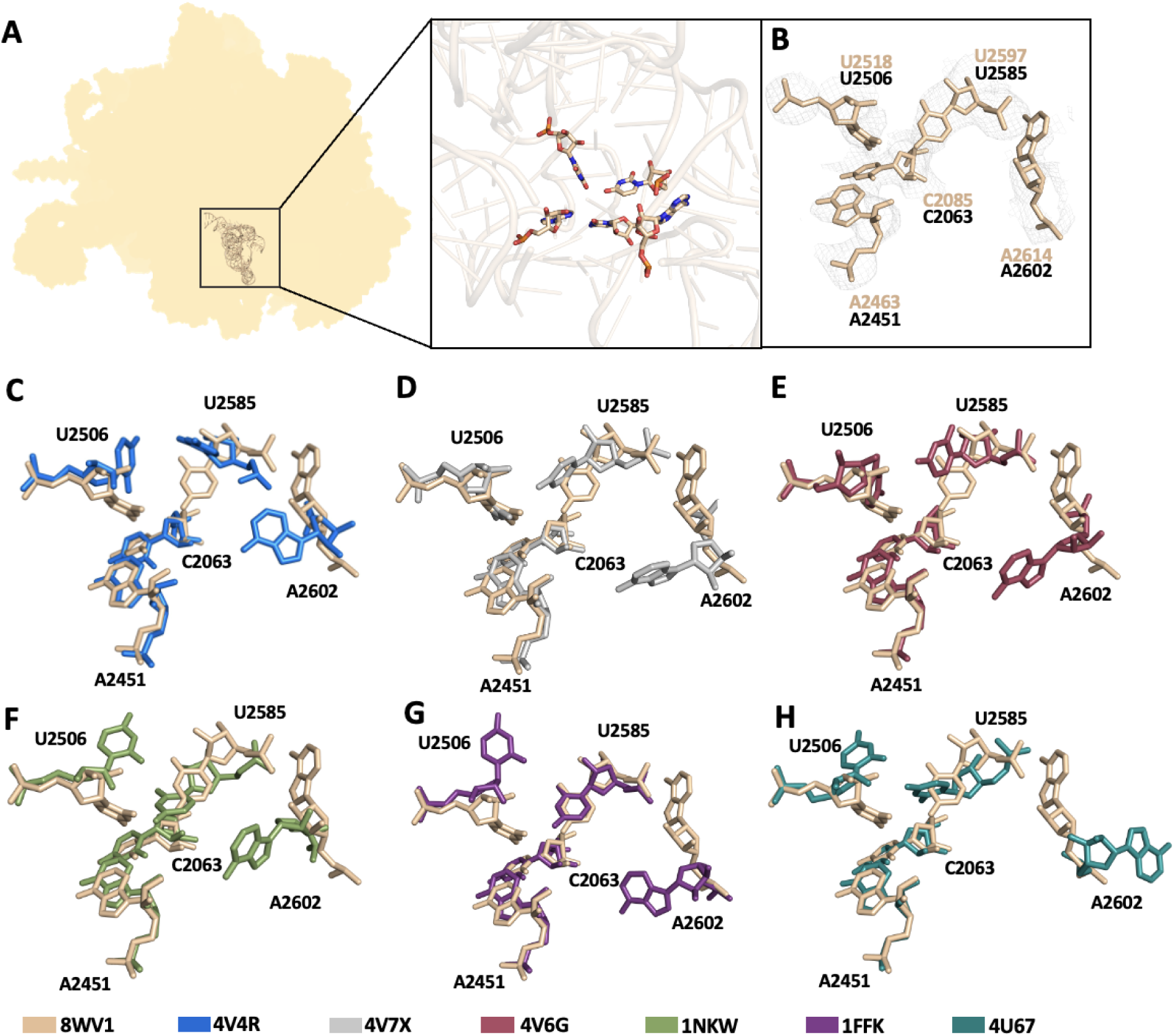
The comparison of evolutionary conserved peptidyl-transferase center residues with the ambient temperature structure of 50S ribosomal subunit from *T. thermophilus*. **(A)** The localization of PTC in the 50S subunit and the close view of the peptidyl-transferase center. **(B)** 5 peptidyl-transferase center residues with *2Fo-Fc* simulated annealing-omit map. Comparison of the ambient temperature 50S structure with cryogenic ribosome structures: **(C)** 4V4R; 70S *T. thermophilus* ribosome in complex with Release factor 1 and 2. **(D)** 4V7X; whole *T. thermophilus* ribosome complexed with erythromycin **(E)** 4V6G; 70S *T. thermophilus* ribosome in complex with two tRNAs and mRNA **(F)** 1NKW; 50S ribosomal subunit structure from *Deinococcus radiodurans*. **(G)** 1FFK; 50S ribosomal subunit structure from *Haloarcula marismortui*. **(H)** 4U67; 50S ribosomal subunit structure from *Deinococcus radiodurans* with a 3 residue insertion in L22.

A2602 plays a pivotal role in the hydrolytic activity of the active site during the release of nascent proteins (Polacek et al., 2003; Youngman et al., 2004). Our study confirmed the intrinsic flexibility of these residues, particularly A2602, at ambient temperature. Some prior crystallographic investigations have indicated that the orientation of these residues is influenced by the binding of peptidyl transferase substrates, with A2602 undergoing the most pronounced conformational changes upon substrate binding (Nissen et al., 2000; Schmeing et al., 2002). However, it remains unclear whether the flexibility of any of these residues holds functional significance for either the peptidyl transfer process or the hydrolysis of peptidyl-tRNA.

### Antibiotic binding to 50S

The effect of temperature on possible interactions and conformational changes upon binding of two major classes of antibiotics was investigated at the nucleotide scale. We superposed our 50S^AT^ structure with 50S^CT^ structures containing erythromycin (PDB: 4V7X) and chloramphenicol (PDB: 4V7W) (Bulkley et al., 2010). Additionally, the apo form of the 50S^CT^ structure was obtained from crystals of the 70S T. *thermophilus* (PDB:4V8I) (Polikanov et al., 2012). The sequential comparisons allowed us to identify drug-specific and temperature-dependent changes.

The macrolide antibiotic erythromycin binds at the entrance to the exit tunnel, preventing the elongation of newly synthesized polypeptides (Schlünzen et al., 2001). The superposed 50S^AT^ the 50S^CT^ containing erythromycin (PDB ID: 4V7X) (Figure 7A-B) shows conformational differences on residues within the binding pocket (Figure 7C-E-G). The residues U2611, A2058, and A2059 are found in a proper orientation to participate in electrostatic interactions with the lactone ring of erythromycin (Figure 7G-H). However, our comparisons revealed that nucleotide A2062 can occupy different states along the three structures. Although the ribose rings within the 50S^AT^ and 50S^CT_Apo^ structures (PDB ID: 4V8I) are oriented similarly, the bases are rotated at the glycosidic bonds. In contrast, the erythromycin-bound 50S^CT_Holo^ structure shows A2062 rotated at the backbone (Figure 7C-D-E-F-G).

**Figure 7.**
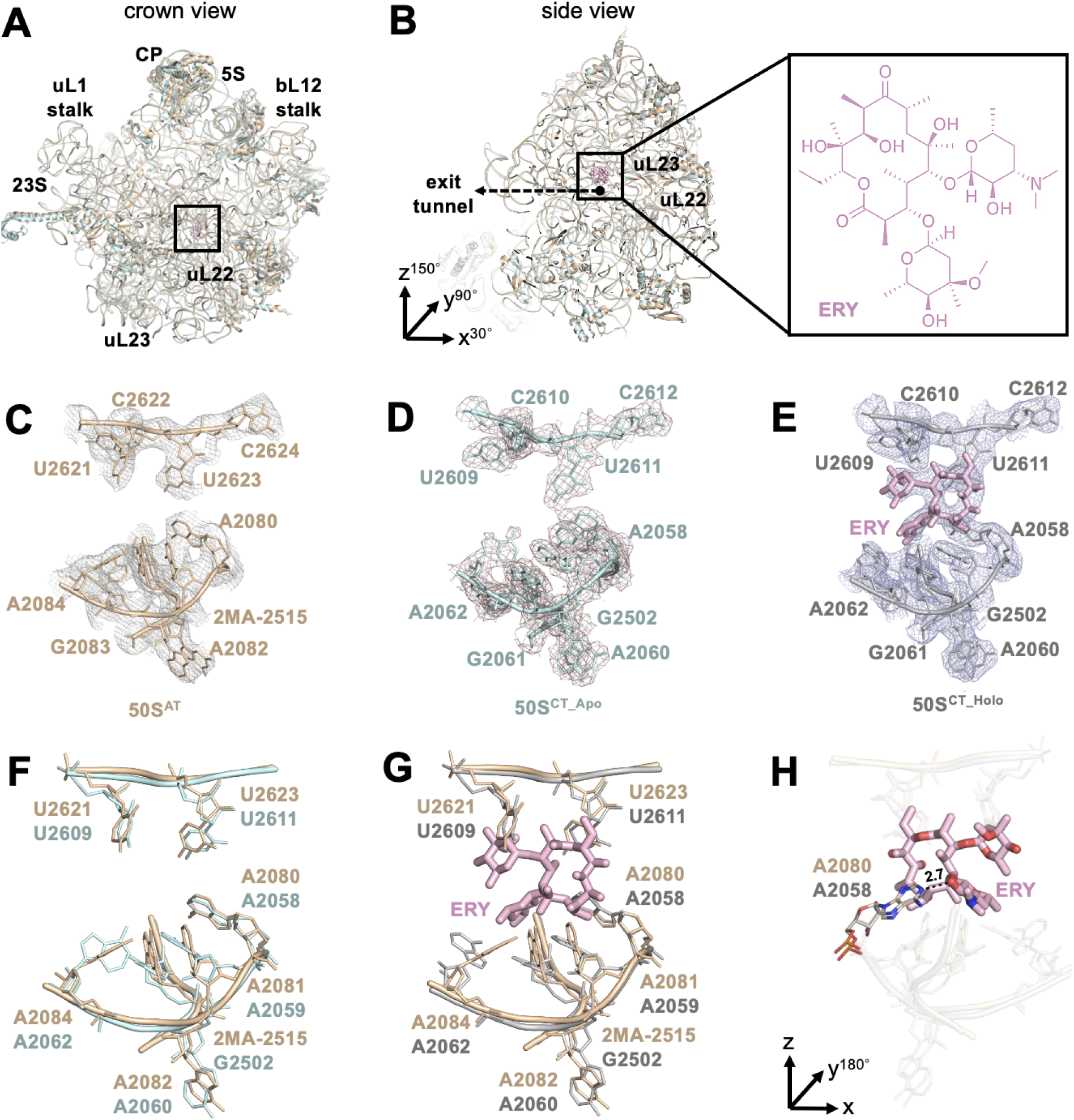
Comparison of the apo 50S structure at ambient temperature, at cryogenic temperature (PDB ID: 4V8I), and the erythromycin (ERY) bound structure (PDB ID: 4V7X) shows minor conformational change in A2062. **(A-B)** 50S ribosomal subunits are superposed and the position of erythromycin is indicated with a square. For panel B, the overall structure is rotated based on coordinates x=30°, y=90° z=150°, respectively. **(C-E)** 2*Fo-Fc* simulated annealing-omit map for 50S structures is shown in gray, dark violet and light blue, respectively and contoured at 1.0 σ level. **(F)** 50S ribosome at ambient temperature 50S^AT^ is superposed with the 50S subunit at cryogenic temperature 50S^CT^ with the RMSD value of 0.858 Å. **(G-H)** 50S subunit is superposed with the erythromycin bound structure with the RMDS value of 0.985 Å. Erythromycin is colored in light pink and hydrogen bond interaction (2.7 Å) with the critical residue A2058 is shown with a dashed black line. To get the view position, the overall structure in panel G is turned 180° in y axis. All residues are labeled based on *E.coli* numbering.

At the peptide exit tunnel, chloramphenicol binds near the peptidyl transferase center and blocks peptide bond formation (Tereshchenkov et al., 2018). Unlike macrolides, chloramphenicol has no lactone ring, limiting its stabilization at the binding site (Figure 8B). In an early study by Bulkley et al., chloramphenicol was found to interact with C2452 of 23S RNA and a potassium cation coordinated near the peptidyl transferase center in the cryogenic 70S *T. thermophilus* ribosome (50S^CT_Holo^) (Bulkley et al., 2010). The potassium cation is not visible in the electron density map of our structure, but the positions of the residues G2061, C2501, and G2447, which coordinate the cation in the 50S^CT_Holo^ structure, remain the same (Figure 8C-D-E-F-G-H). In addition to stabilization through the potassium ion, chloramphenicol forms a pi-stacking interaction with the 23S rRNA residue C2452 by its nitrobenzyl ring (Bulkley et al., 2010). The superposition of the 50S^AT^ with the chloramphenicol-bound 50S^CT^ structure revealed a slightly different conformation at the backbone of C2452 depending on temperature (Supplementary Figure 5). On the other hand, the adjacent residue A2453 showed flexibility due to temperature by flipping slightly to the opposite side (Supplementary Figure 5). These findings emphasize the importance of temperature on the conformation of critical residues for both antibiotic binding and ribosomal functions.

**Figure 8.**
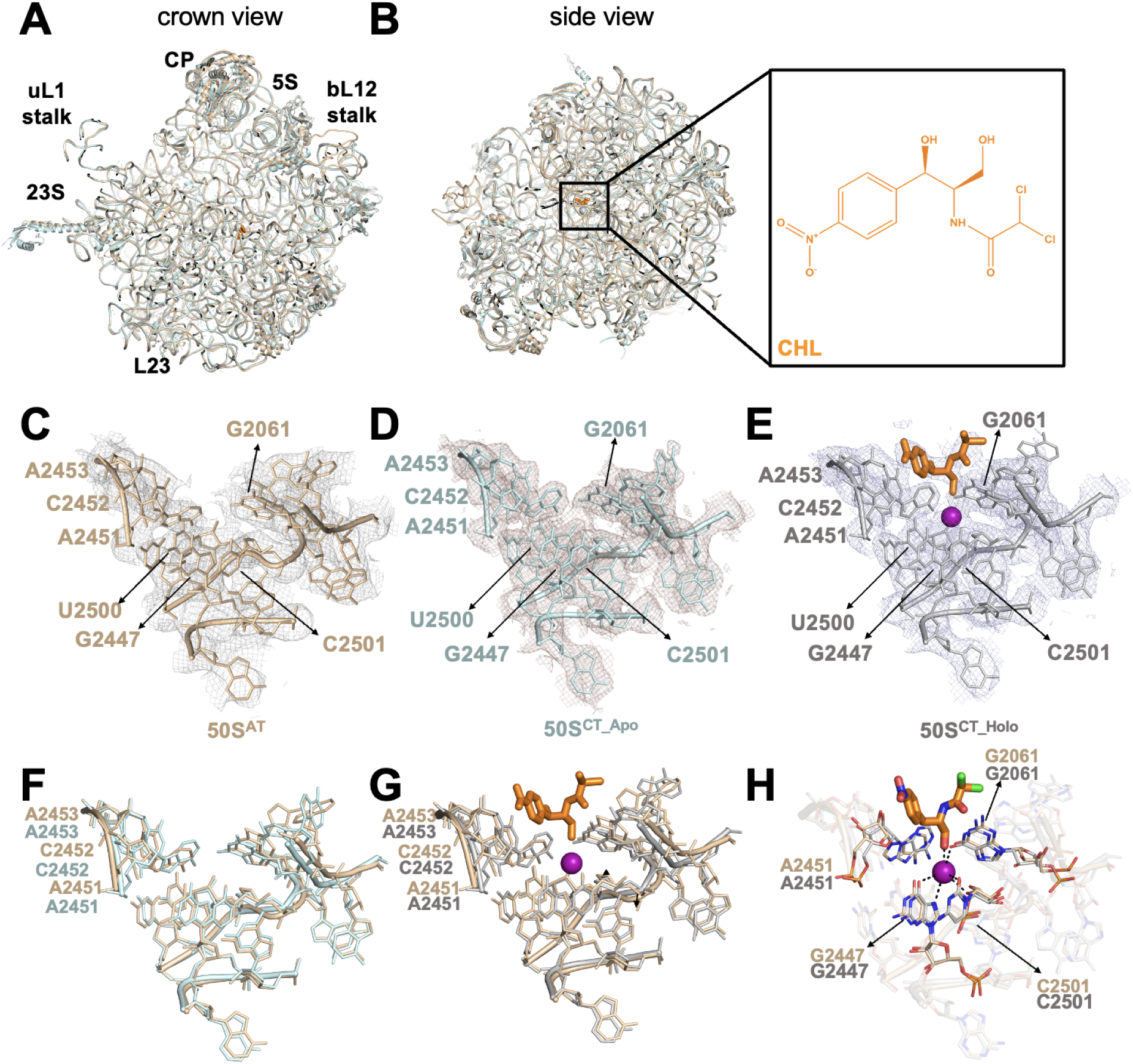
Comparison of the apo 50S^AT^ with the 70S^CT^ structure (PDB ID: 4V8I) and the chloramphenicol (CHL) bound structure (PDB ID: 4V7W) shows minor conformational changes in C2452 and A2453. **(A-B)** 50S ribosomal subunits are superposed and the position of chloramphenicol is indicated with a square. For panel B, overall structure is rotated based on coordinates x=30°, y=270° z=180°, respectively. **(C-E)** *2Fo-Fc* simulated annealing-omit map for 50S structures is shown in gray, dark violet and light blue respectively for ambient, cryo-apo and cryo-antibiotic bound, and contoured at 1.0 σ level. **(F)** The 50S^AT^ is superposed with the 50S subunit at cryogenic temperature with the RMSD value of 0.657. **(G-H)** The 50S ribosome superposed with the chloramphenicol bound structure with the RMDS value of 0.64. Chloramphenicol is colored in orange and hydrogen bond interactions of A2451, G2447, C2501, and G2061 are shown with dashed lines.

## DISCUSSION

Here we present the structure of a dimeric complex of the *T. thermophilus* 50S subunit using XFEL data obtained at ambient temperature to a resolution of 3.99 Å (Figure 1). This work presents a streamlined method for preparing *T. thermophilus* 50S ribosomal subunit microcrystal samples with volumes less than 1 mL, which is sufficient to produce a full preliminary ambient temperature structure at an XFEL light source. The diffraction data collected at the Linear Coherent Light Source (LCLS) resulted in 1000 indexed patterns in 47 minutes of data collection, while also consuming approximately 50 uL of crystal sample slurry. This structure highlights the potential for SFX experiments to produce structures that would be difficult or impossible to obtain using synchrotron X-rays due to the high radiation sensitivity of a very large asymmetric unit cell with limited lattice contacts. The dimeric form of the *T. thermophilus* 50S subunit is one of the largest biological macromolecule structures determined using the SFX technique at an XFEL to date. This dimerization was first observed in some of the earliest electron microscopy experiments (Huxley & Zubay, 1960). Since information about the dimeric form of 50S ribosomal subunits in a crystal structure is very limited in the literature, this structure provides valuable information about dimerization of the large ribosomal subunit.

Magnesium ions can be found stabilizing many structural elements within the ribosome as described in Klein et al. (Klein et al., 2004). They neutralize the tight packing of negatively charged rRNA, displacing associated water molecules and result in specific binding (Klein et al., 2004). Compared with other ribosome structures, our 50S dimer contains a large number of hexahydrated magnesium ions, likely due to collection by the diffract-before-destroy approach that avoids the extensive radiation damage expected by other techniques. Both hydrated and dehydrated magnesium cations were identified based on their electron density and computational methods; this expansion in solvent information can be of benefit to future computational studies aimed at identifying magnesium ions important for structural stability.

To shed light on the ambient temperature conformations of ribosomal proteins, a GNM analysis was performed for the evolutionary conserved protein uL23 located near the peptide exit tunnel. The protein has been associated with the folding of the nascent polypeptide chain and assisting the movement of newly synthesized polypeptide chains through the peptide exit tunnel (Gloge et al., 2014). The unexpected difference in theoretical and experimental mobility of the stem emphasizes the importance of the temperature effect on the protein conformational stability. The loop region of both complexed and uncomplexed uL23 showed mobility. While the movement of ribosome-assembled uL23 is restricted to the tip of the loop due to the salt bridge between uL23 Arg68 and 23S rRNA C482, the cumulative 10 slowest modes revealed that it acts as a hinge in low-frequency motions.

A better understanding of the dynamics of the 50S subunit opens up new possibilities for drug design approaches. Conformational changes caused by antibiotics are expected to play a critical role in the mechanism of inhibition. Significant conformational changes are revealed near the Erythromycin binding site by the comparison of cryogenic apo and erythromycin-bound structures. Minor conformational changes are detected for the residues U2062, U2060 and A2059, while minor changes are observed for A2058. As A2058 is a critical residue for drug binding, these data may suggest that it exhibits higher stability near the exit tunnel compared to neighboring residues.

The comparative analysis of the ambient temperature structure with the apo and bound cryo-structures revealed the conformational dynamics of the chloramphenicol binding site. In the cryogenic 50S^CT_Holo^ structure, the residues G2061, C2501, and G2447 coordinate a potassium. Temperature does not appear to be affecting the conformation of the potassium coordinating residues, but slightly different conformations have been observed for residues in and around the peptidyl-transferase center. Previously it has been stated that peptidyl-transferase activity is dependent on both the presence of the K^+^ ion and temperature (Miskin et al., 1968; Vogel et al., 1971; Rodriguez-Correa & Dahlberg, 2008). C2452 is a critical residue for chloramphenicol binding since it stabilizes the *p*-nitrobenzyl group of the antibiotic, and this residue is found to be more flexible in its backbone at ambient temperature in our structure.

Determining the actual conformation of the residues at ambient temperature becomes more crucial when the main driving force of the enzymatic activity is considered. Previous studies showed that the peptidyl-transfer mechanism is entropic rather than enthalpic in nature. Most enzymes catalyze their reaction enthalpically by stabilizing their substrates with polar interactions to lower activation energy. However, the peptidyl-transferase center of the ribosome acts as an entropy trap for the catalysis of peptide bonds (Sievers et al., 2004; Gregory & Dahlberg, 2004). The active site assists bond formation by positioning the substrates in the correct orientation and preserving the catalytic environment from water molecules. In the ambient temperature structure, A2453 demonstrated a different conformation, revealing the more precise orientation and flexibility for peptide bond formation.

This study provides valuable information that could be useful in addressing the issue of antibiotic resistance, which has become a greater concern. Understanding the structural dynamics of the apo and antibiotic-bound 50S ribosomal subunit may pave the way for the development of new generation antibiotics that can target the protein synthesis machinery more selectively and effectively. Besides its contribution to antibiotic studies, these results are important in the application of XFEL technique in ribosome studies, and lay the foundations for structural studies of ribosomes at ambient temperature.

## DATA AVAILABILITY

The structure and its electron density map determined and analyzed in this study have been deposited in Protein Data Bank (PDB) under accession number 8WV1.

## SUPPLEMENTARY DATA

Supplementary Data are available at NAR online.

## ACKNOWLEDGEMENT

Authors would like to dedicate this manuscript to the memory of Dr. Albert E. Dahlberg and Dr. Nizar Turker. We thank the staff at Linear Coherent Light Source at SLAC National Accelerator Laboratory for their assistance in data collection. We thank the Photon Ultrafast Laser Science and Engineering Institute (PULSE Institute) at Stanford University for providing technical support.

## FUNDING

Use of the Linac Coherent Light Source (LCLS), SLAC National Accelerator Laboratory, is supported by the U.S. Department of Energy, Office of Science, Office of Basic Energy Sciences under contract no. DE-AC02-76SF00515. H.D. acknowledges support from NSF Science and Technology Center grant NSF-1231306 (Biology with X-ray Lasers, BioXFEL). This publication has been produced benefiting from the 2232 International Fellowship for Outstanding Researchers Program, 2236 CoCirculation2 program, the 1001 Scientific and Technological Research Project Funding Program and the 2244 Industry Academia Partnership Research Project Funding Program of the TÜBİTAK (Project Nos. 118C270, 121C063, 120Z520 and 119C132). However, the entire responsibility of the publication belongs to the authors of the publication. The financial support received from TÜBİTAK does not mean that the content of the publication is approved in a scientific sense by TÜBİTAK.

SLAC National Accelerator Laboratory; access to the Linac Coherent Light Source CXI instrument.

## Conflict of interest statement

None declared

## Notes

### Competing Interest Statement

The authors have declared no competing interest.

