## Supplementary File for "Ambient Temperature Bacterial Large Ribosomal Subunit Structure Enabled by Serial Femtosecond X-ray Crystallography"

**Supplementary Table 1:** X-ray data collection and refinement statistics

| ***Thermus thermophilus* 50S ribosomal subunit (XFEL)** | |
|---|---|
| *Data collection* | PDB ID: 8WV1 |
| Beamline | LCLS (CXI) |
| Space group | $P4_1$ |
| **Cell dimensions** | |
| *a*, *b*, *c* (Å) | 303.8, 303.8, 434.8 |
| α, β, γ (°) | 90, 90, 90 |
| *R*split | 141.29% |
| Completeness (%) | 0.56 @ 3.9, 0.78 @ 3.4, 0.9 @ 4.56 |
| SFX multiplicity of observations | |
| *CC** | 0.8633217 |
| Wilson B Factor (Å2) | 51.80 |
| **Refinement** | |
| Resolution (Å) | 3.99 |
| No. of reflections | 286015 |
| *R*work/*R*free | 0.2957/0.3617 |
| Ramachandran favoured (%) | 96.88% |
| Ramachandran allowed (%) | 3.07% |
| Ramachandran outliers (%) | 0.05% |
| **No. of atoms** | |
| Protein | 31250 |
| RNA | 61845 |
| Other | 14060 |
| ***B*-factors (Å2)** | |
| Protein | 73.40 |
| RNA | 92.81 |
| Other | 50.65 |
| **Root-mean-square deviations** | |
| Bond lengths (Å) | 0.002 |
| Bond angles (°) | 0.521 |

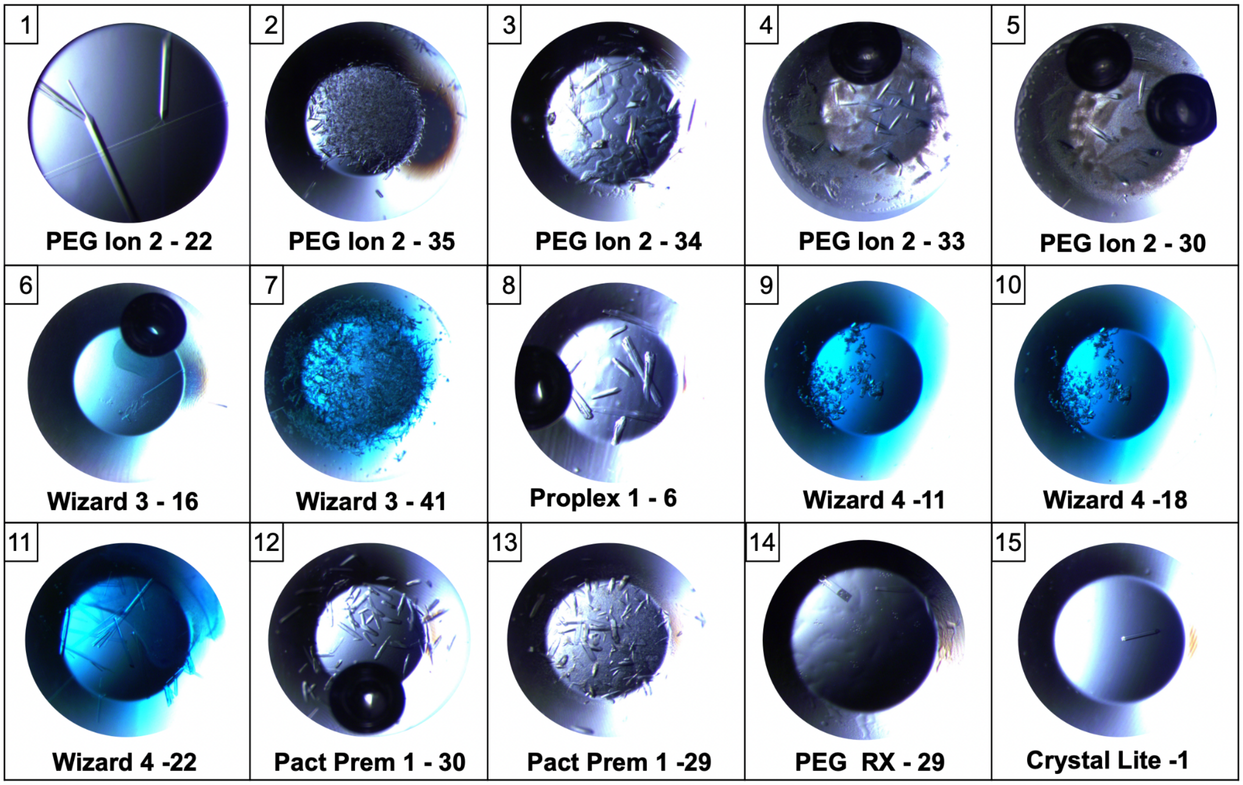


**Supplementary Figure 1.** The picture of 50S ribosomal subunit under light microscope. (1) First crystal picture represents the Synchrotron Crystal approximately 9Å resolution. (2) Second crystal picture represents SFX crystals beyond 4Å resolution.

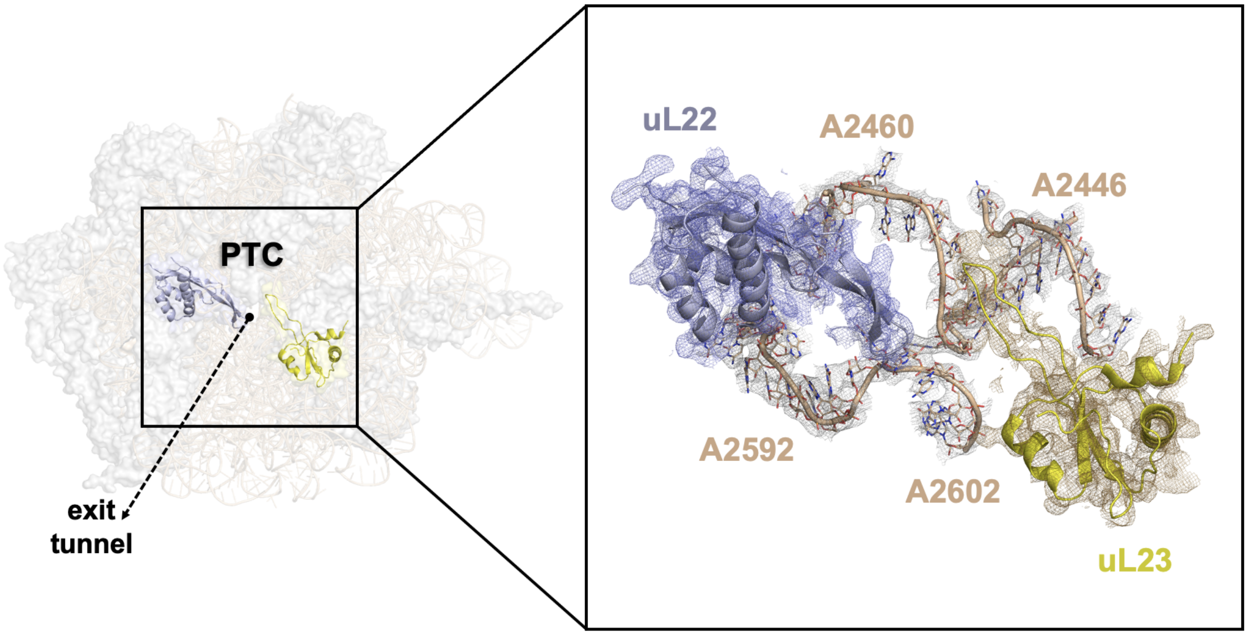

**Supplementary Figure 2.** Representation of well-defined electron density map near to the peptidyl transferase center (PTC). 2Fo-Fc electron density map is colored based on each structure and contoured at the 1σ level.

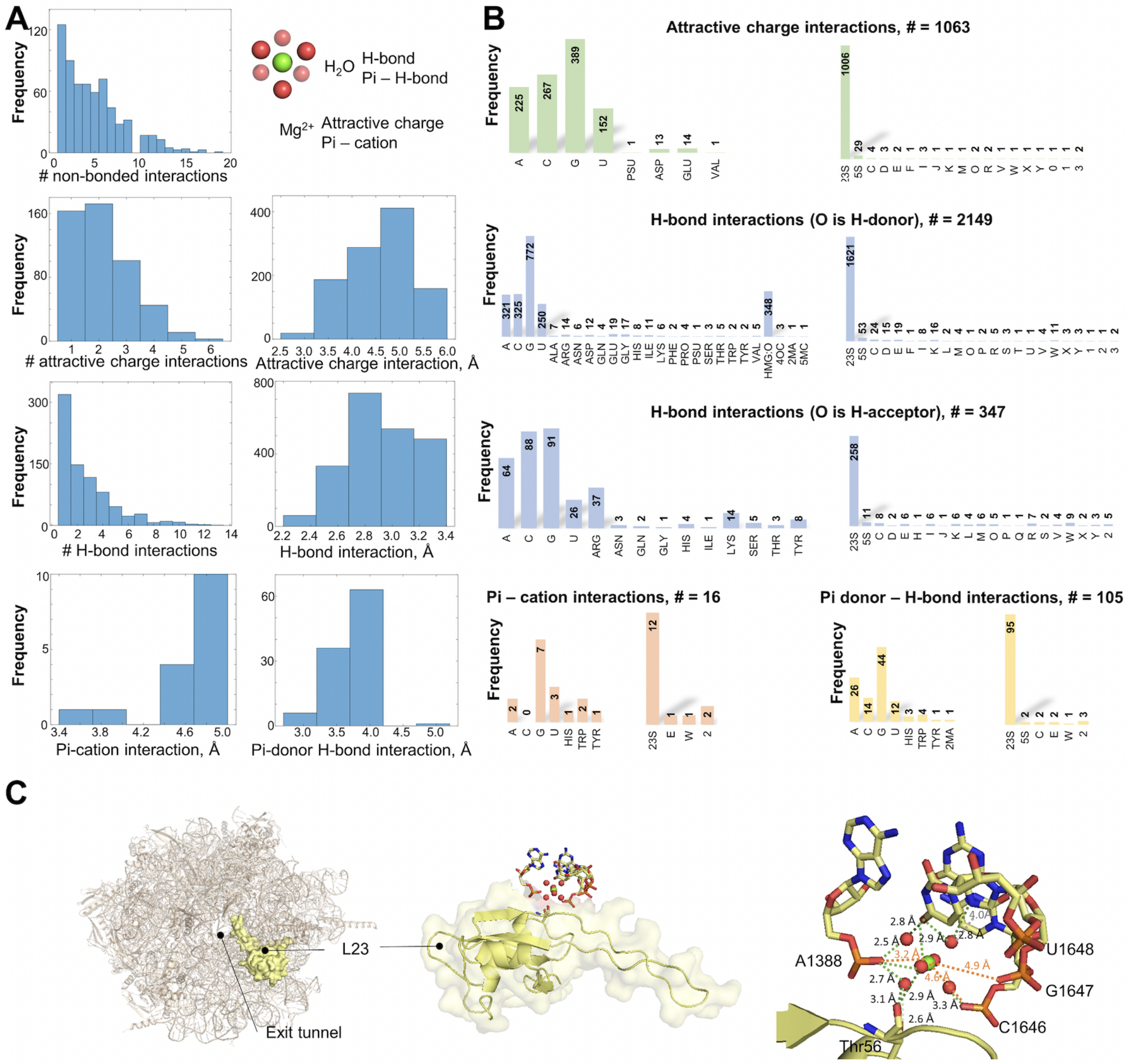


**Supplementary Figure 3. (A)** Histograms for different types of non-bonded interactions of hexahydrated magnesium groups on one 50S subunit. Magnesium ions make attractive charge and pi-cation interactions, while oxygen atoms establish H-bond and pi donor – H-bond interactions. **(B)** Histograms for non-bonded interactions involving different residues and different chains named as in the structural data. **(C)** Non-bonded interactions between the hexahydrated magnesium group at the interface of 23S rRNA and ribosomal protein L23.

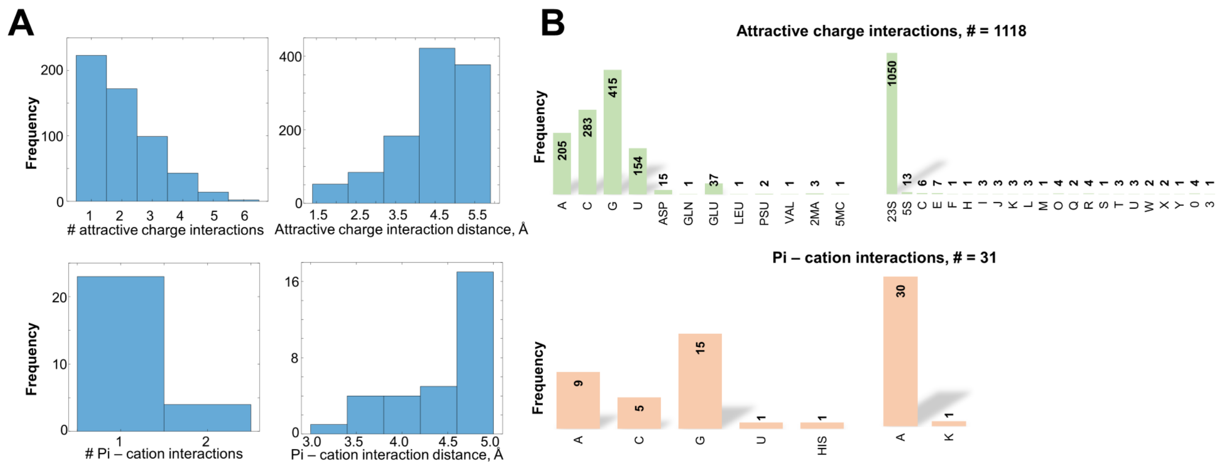


**Supplementary Figure 4. (A)** Histograms for different types of non-bonded interactions of magnesium ions on one 50S subunit. Magnesium ions make attractive charge and pi-cation interactions. **(B)** Histograms for non-bonded interactions involving different residues and different chains named as in the structural data.

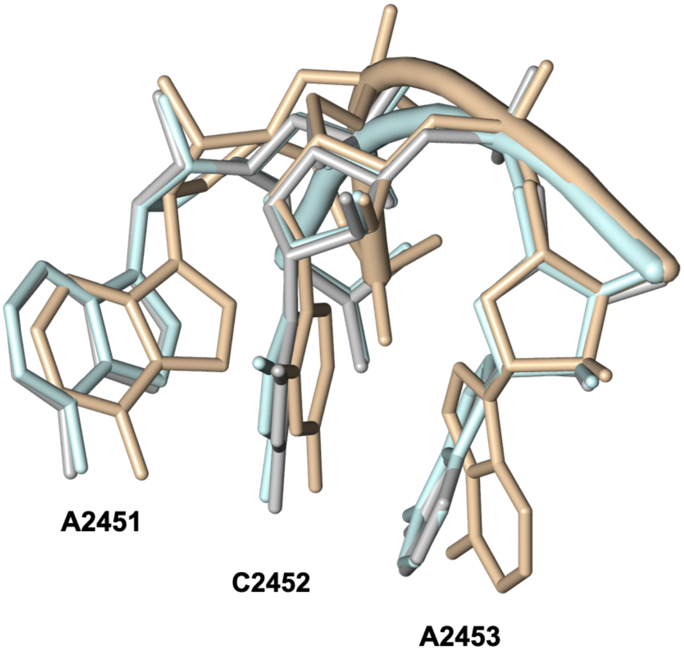

**Supplementary Figure 5.** Nucleotide residues A2451, C2452, A2453 located near the chloramphenicol binding site. The residues of the ambient 50S structure are colored with wheat, apo cryogenic 50S colored with pale cyan, chloramphenicol bound cryogenic 50S colored with gray. The figure demonstrates the conformational change of the residues due to the change in temperature. A conformational change at the A2453 is found to be remarkable.
